# COPS5 is Essential for Sertoli Cell Function and Male Fertility in Mice

**DOI:** 10.64898/2025.12.19.695357

**Authors:** Changmin Niu, Tao Li, Wei Li, Yi Tian Yap, Qian Huang, Lei Jiang, Opeyemi Dhikhirullahi, Ava Miciuda, Eva Faddoul, Shizheng Song, Michael D Griswold, Zhibing Zhang

## Abstract

The COP9 signalosome subunit 5 (COPS5) is a multifunctional protein that regulates ubiquitin-dependent processes. Global knockout of *Cops5* is embryonically lethal, and while it is known to be vital in germ cells and testicular smooth muscle, its function in Sertoli cells, the key somatic supporters of spermatogenesis, remains entirely unknown. This study investigates the critical role of COPS5 within Sertoli cells. Using Sertoli cell-specific *Cops5* knockout mouse models, we demonstrate that COPS5 is essential for maintaining male fertility. Sertoli-specific *Cops5* ablation resulted in age-dependent male infertility, despite normal initial development. Mutants exhibited progressive testicular atrophy, oligoasthenospermia, and significantly reduced testis weight. Histology showed vacuolated, disorganized tubules devoid of germ cells and sperm. Crucially, COPS5 loss disrupted Sertoli cell polarity, evidenced by aberrant cytoplasmic mislocalization of the nuclear marker WT1 and detachment from the basement membrane. Integrity of the blood-testis barrier (BTB) was severely compromised, with discontinuous and punctate expression of the tight junction adaptor ZO-1. Intercellular communication was also impaired, shown by a stark reduction in Connexin 43 gap junction signals. Our findings establish that COPS5 is an indispensable intrinsic regulator of Sertoli cell function. Its loss disrupts cell polarity, BTB architecture, and gap junction communication, leading to failed support of spermatogenesis and consequently infertility. This work defines a novel and critical somatic function for COPS5 in male reproduction. We conclude that COPS5 is intrinsically required in Sertoli cells to maintain their polarity and BTB function, which are foundational for supporting germ cell development and ensuring male fertility. This identifies COPS5 as a novel, essential regulator within the testicular somatic compartment.

## Introduction

Infertility has become a public health problem globally, with recent epidemiological studies indicating that it affects approximately 17.5% of the adult population, with male factors being a sole or contributing cause in roughly 50% of cases ^[1,2]^. Spermatogenesis, the complex and highly regulated process of sperm production, is fundamental to male fertility. This process occurs within the seminiferous tubules of the testis and relies not only on germ cells but also on intricate somatic cell support. Sertoli cells, the epithelial cells of the tubules, provide structural and nutritional support and form the blood-testis barrier. Leydig cells, located in the interstitium, are responsible for producing testosterone, a hormone indispensable for the completion of meiosis and spermiogenesis ^[3–5]^. The precise coordination between these somatic cells and the developing germ cells is crucial for producing functional sperm. Disruptions in this delicate cellular crosstalk are a significant cause of idiopathic male infertility, highlighting the critical importance of identifying and characterizing key regulatory genes within the testicular somatic compartment ^[6–7]^.

The COP9 signalosome complex subunit 5 (COPS5), also known as CSN5 or Jab1, is the catalytic core of the conserved eight-subunit COP9 signalosome (CSN) complex ^[8]^. Structurally, COPS5 contains an N-terminal MPN (Mpr1-Pad1-N-terminal) domain that confers its isopeptidase activity, which is essential for the deneddylation of cullins within Cullin-RING E3 ubiquitin ligases (CRLs) ^[9]^. This activity is paramount for maintaining the specificity and cycling of CRLs, thereby regulating the ubiquitin-mediated degradation of a vast array of substrate proteins involved in diverse cellular processes ^[10–11]^. COPS5 is ubiquitously expressed in mammalian tissues, with the highest expression level in testicular tissue and is predominantly localized in the nucleus, though it can shuttle to the cytoplasm ^[12–13]^. Beyond its core function in the CSN complex, COPS5 has been reported to interact independently with numerous client proteins, such as p53, c-Jun, and HIF-1α, positioning it as a central regulator of critical biological processes including cell cycle progression, DNA damage response, apoptosis, and oncogenesis ^[9,14–16]^.

Global deletion of *Cops5* in mice results in early embryonic lethality due to severe developmental defects, alluding to its essential role in early embryonic development and growth potential ^[8]^. This lethality has necessitated the use of tissue-specific knockout models to unravel its postnatal roles. For instance, conditional knockout of Jab1 in mouse limb buds and chondrocytes results in severely shortened limbs and neonatal lethal chondrodysplasia, respectively ^[17]^. Specific knockout of *Cops5* in osteoblast precursor cells exhibited the essential role of COPS5 in proper bone growth and survival ^[9,17–18]^, while its deletion in germ cells exhibited meiotic arrest and complete infertility, demonstrating its essential role in maintaining male germ cell survival and acrosome biogenesis ^[19]^. Moreover, a recent study employing a *Myh11*-Cre driver to delete *Cops5* in smooth muscle cells demonstrated its essential role in spermatogenesis, with knockout males exhibiting developmental disorders, failed development of reproductive organs, impaired endocrine system associated with testicular functions, and spermatogenesis defects ^[20]^. However, germ cells and smooth muscle cells’ specific ablation leave a fundamental question unanswered: what is the functional requirement for COPS5 within the other testicular somatic cells, such as Sertoli cells and Leydig cells? The function of COPS5 in somatic cells remains entirely unexplored.

Sertoli cells are the central somatic custodians of spermatogenesis ^[21]^. Their highly polarized and intricate cytoplasmic structure forms the specific microenvironment of the seminiferous epithelium, which is essential for germ cell development. A defining functional feature of Sertoli cells is the formation of the blood-testis barrier (BTB), which creates an immunologically privileged compartment for meiosis and post-meiotic development. Beyond this structural role, Sertoli cells are metabolically active, providing nutrients and growth factors to germ cells, phagocytosing residual bodies, and secreting fluid essential for spermiation ^[22–26]^. The absolute necessity of Sertoli cells is evidenced by the fact that the ablation of any key regulatory gene specifically within this cell type, such as those involved in androgen signaling, retinoic acid metabolism, or cytoskeletal organization-invariably, leads to a failure in spermatogenesis and consequently male infertility ^[27–30]^. Their functional integrity is therefore the cornerstone of male fertility.

Given the established embryonic lethality of the global knockout, the recent discovery of its crucial role in testes, there is a compelling and unmet need to investigate the role of COPS5 specifically in Sertoli cells. Therefore, this study is designed to definitively address this question by generating and characterizing Sertoli cell-specific *Cops5* conditional knockout mouse model (*Cops5* ^fl/fl^; *Amhr2*-Cre^/+^). We demonstrate that COPS5 is intrinsically required for Sertoli cell function and that its ablation will disrupt the BTB integrity, Sertoli cell polarity and spermatogenesis, leading to male infertility. Our findings will elucidate a novel and essential somatic function for COPS5 in male reproduction.

## Materials and Methods

### Ethics statement of animal use

All mouse research was approved by Wayne State University Institutional Animal Care with the Program Advisory Committee (Protocol number: 24-02-6561).

### Generation of Sertoli cell-specific Cops5 knockout mice

The *Cops5* ^flox/flox^ (*Cops5* ^fl/fl^) mice were kindly provided by Dr. Ruggero Pardi (Università Vita, Italy). 3-to-4-month-old *Cops5* ^fl/fl^ females were crossed with age-paired *Amhr2-*cre/^+^ males (MMRRC, Stock Number: 014245-UNC) in which the Cre recombinase was driven by the anti-Mullerian hormone receptor type 2 (Amhr2) promoter which directs the expression of Cre recombinase mainly in the Sertoli cells. The resulting *Cops5* ^fl/+^; *Amhr2*-Cre/^+^ males were crossed back with the *Cops5* ^fl/fl^ females. The *Cops5* ^fl/fl^; *Amhr2*-Cre/^+^ were considered to be the Sertoli cell-specific *Cops5* knockout mice (hereafter referred to as *Cops5* cKO). The *Cops5* ^fl/+^; *Amhr2-*Cre/^+^or *Cops5* ^fl/fl^ mice were used as the negative controls. All mice used in this report were on a C57BL/6J background. Mice genotypes were identified by PCR using the following primers:

*Cops5* forward: 5’ -GCCTGCATTACCGGTCGATGCAACGA- 3’;

*Cops5* reverse: 5’ -GTGGCAGATGGCGCGGCA- 3’;

*Amhr2-cre* forward: 5′- CCGCTTCCTCGTGCTTTACGGTAT -3′;

*Amhr2-cre* reverse: 5′- ACCTAGTAGAGAGGCTGCGTTGAGTGTG -3′.

### H&E staining

Testicular and epididymal tissues from control and *Cops5* cKO mice of different ages were fixed in Bouin’s solution (SIGMA, Saint Louis, MO, USA, HT10132), dehydrated through a graded alcohol series, cleared in xylene, and embedded in paraffin. The paraffin blocks were sectioned into 4μm thick slices and mounted onto glass slides for subsequent staining. The sections were deparaffinized in xylene, rehydrated through a graded alcohol series, and stained with hematoxylin and eosin (H&E). After mounting with neutral resin, images were captured under a microscope for histological analysis. The diameter and cross-sectional area of seminiferous tubules were measured using Image J (National Institutes of Health, USA).

### Male fertility assessment

To evaluate male fertility, each 6-week-old to 4-month-old control and *Cops5* cKO mouse was co-caged with a single age-matched, reproductively competent wild-type female for a minimum period of one month. The litter size and offspring viability were subsequently observed and documented, with a sample size of six or more mice (n ≥ 6) per group.

### Sperm parameter analyses

Epididymal cauda were isolated from *Cops5* cKO and control mice at different weeks of age. The tissues were minced into small fragments in human tubal fluid (HTF) solution (Cosmo Bio, Tokyo, Japan, CSR-R-B071) containing 10% FBS (Gibco, California, USA, A5256701) and maintained at 34°C, followed by 7 minutes of incubation. A hemocytometer was used to determine sperm concentration. Sperm motility and kinematic parameters were evaluated using a Nikon TE200E inverted microscope on a prewarmed slide with Sanyo color charge-coupled device, Hi-Resolution camera (VCC-3972) and Pinnacle Studio HD (version 14.0) software. After centrifugation, the sperm pellets were washed with PBS (Gibco, California, USA, 10010023) and fixed in 4% paraformaldehyde (Sigma, California, USA, YB36314ES60) at room temperature for 30 minutes. The fixed sperm were then prepared as smears on microscope slides for subsequent morphological evaluation.

### Immunofluorescence staining

Testicular tissues from mice were fixed in 4% paraformaldehyde for a duration of 48 hours. The fixed tissues were then processed through a series of steps including dehydration in a graded ethanol series clearing, and infiltration with paraffin wax to generate paraffin-embedded blocks. Sections were cut from these blocks at a thickness of 4μm and baked onto slides at 60°C for 4 hours in preparation for immunofluorescence staining. For the staining procedure, the paraffin sections were first deparaffinized and rehydrated through a descending graded alcohol series. To eliminate endogenous peroxidase activity, the sections were treated with a solution of methanol and hydrogen peroxide in a 37°C water bath for 10 minutes. After washing thoroughly with ddH_2_O, heat-induced antigen retrieval was conducted using a citrate-based buffer solution. Non-specific binding sites were blocked by incubating the tissues with 1% donkey serum in PBS at room temperature for 1.5 hours. The sections were subsequently incubated with the primary antibody overnight at 4°C. On the following day, the slides were equilibrated to 37°C for 30 minutes in an incubator. Following a wash with PBS, the sections were incubated with fluorescent-labeled secondary antibodies for 2 hours at room temperature under light-protected conditions. Sections were then washed and mounted using VectaMount with DAPI (Vector Laboratories, Burlingame, USA). Images were captured using a Zeiss LSM 700 fluorescence microscope.

The antibodies used in this study are as follows: Sertoli cell marker WT1 (Proteintech, 12609-1-AP, Dilution ratio: 1:200); germ cell marker DDX4 (Abcam, Cambridge, UK, ab27591, Dilution ratio:1:400); Tight junction Adapters protein ZO-1 (Thermofisher, Carlsbad, CA, USA 33-9100, Dilution ratio: 1:200); Gap junction Integral membrane protein Connexin 43 (Cell Signaling Technology, Danvers, MA, USA, 3512S, Dilution ratio: 1:200); Goat Anti-Mouse IgG H and L (Abcam, Cambridge, UK, ab150113, Dilution ratio: 1:2000); Goat Anti-Rabbit IgG H and L (Abcam, Cambridge, UK, ab150078, Dilution ratio: 1:2000).

### Statistical analysis

Data is presented as the mean ± standard error of the mean (SEM). Statistical analysis was performed using Student’s t-tests and Mann-Whitney U tests between groups, where P < 0.05 was considered significant.

## Results

### Generation of Sertoli cell-specific *Cops5* knockout mouse model

AMHR2, the receptor for AMH, exhibits a slight delay in its expression onset compared to the ligand itself. Its expression begins after Sertoli cells have differentiated in the embryonic period and is localized to the already differentiated Sertoli cells. *Amhr2*-Cre is a commonly used tool for achieving Sertoli cell-specific gene knockout in males. To explore the role of COPS5 in Sertoli cells, we generated Sertoli cell-specific *Cops5* conditional knockout mice (*Cops5* ^fl/fl^; *Amhr2*-Cre/^+^) by mating *Cops5* ^fl/fl^ females with *Amhr2*-cre/^+^ male mice. Genotypic identification was used to verify the successful construction of the *Cops5* mutant mouse model. *Cops5* ^fl/+^; *Amhr2-*Cre/^+^ and *Cops5* ^fl/fl^ mice were used as the negative controls (**Supplemental Figure 1**).

### Disruption of *Cops5* gene in Sertoli cells results in progressively reduced male fertility

The *Cops5* cKO mice were grossly normal in both males and females. To test fertility of the cKO males, 6-week-old, two-month-old, three-month-old and four-month-old *Cops5* control and cKO males were bred with wild-type females for one month. All 6-week-old and 2-month-old control and cKO mice showed normal fertility. All breeding mice sired pups, and no significant difference was found in fertility rate and litter size between the control and cKO mice. However, the 3-month-old cKO mice showed significantly reduced fertility rate and litter size. 4-month-old cKO mice completely lost fertility (**Figure 1**).

**Figure 1.**
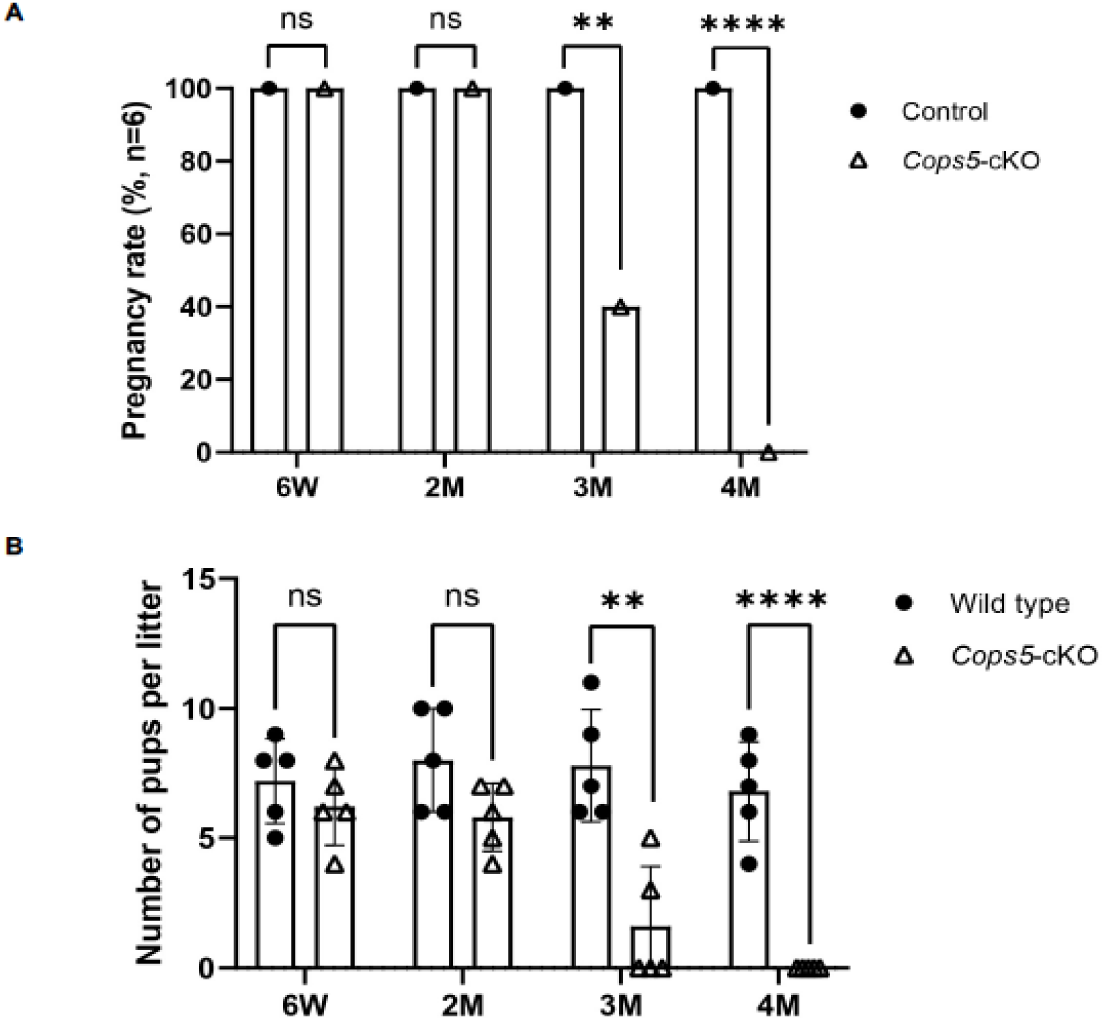
Progressively reduced male fertility in the Sertoli-cell specific *Cops5* cKO mice. The control and cKO mice at the indicated ages were bred with wild-type females for one month, The fertile males and the litter size were recorded. A. Fertility rate of control and cKO mice. No significant difference was found between the control and cKO at 6-week-old and 2-month-old mice. The fertility rate was significantly lower in the 3-month-old cKO mice. The cKO mice lost fertility at 4-month-old and after. n=5. *P<0.05 compared to the control group. B. Litter size of control and cKO mice. No significant difference was found between the control and cKO at 6-week-old and 2-month-old mice. The litter size was significantly lower in the 3-month-old cKO mice. The cKO mice lost fertility at 4-month-old and after. n=5. *P<0.05 compared to the control group.

### The absence of COPS5 in Sertoli cells results in progressively reduced sperm number and motility

To determine the potential factors underlining the progressively reduced fertility, we analyzed sperm parameters in control and cKO mice at various ages corresponding to the fertility study. Sperm were collected from the cauda epididymis of these mice and sperm counts, morphology and motility were examined. There was no significant difference between the control and cKO in these parameters when 6-week-old and 2-month-old mice were analyzed. The 3-month-old *Cops5* cKO mice displayed a pronounced decrease in epididymal sperm number and a significant drop in the percentage of morphologically normal sperm. Sperm motility was also significantly reduced compared to the 3-month-old control mice. By 5 months of age, only a scarce number of sperm could be found in the epididymis of the cKO mice, and virtually none had a normal morphology. The few sperm discovered completely lost motility (**Figure 2**). Collectively, deletion of *Cops5* in Sertoli cells triggered progressive defects in sperm production and function.

**Figure 2.**
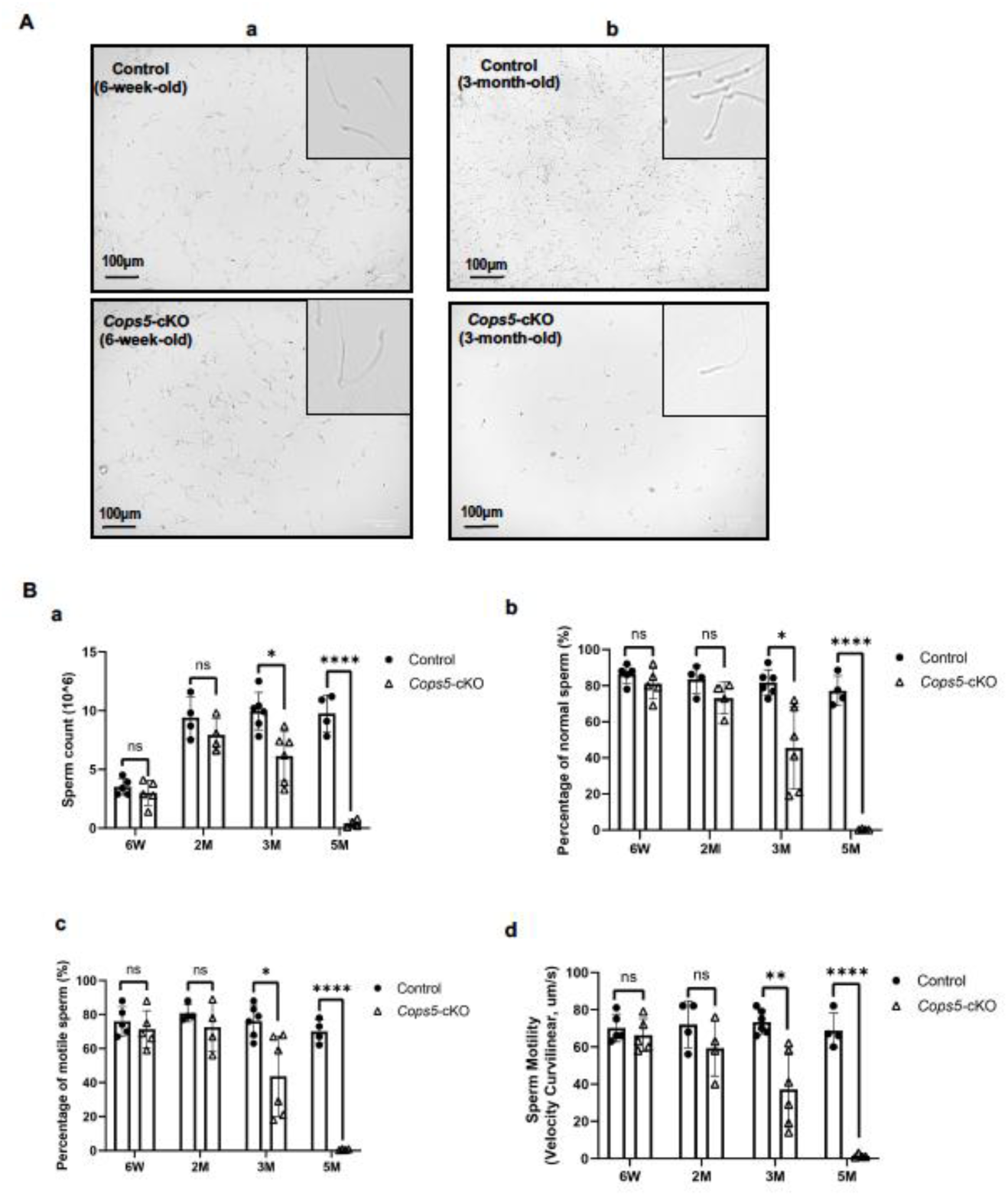
Progressively reduced sperm number and motility in the cKO mice. Sperm were collected from cauda epididymis and morphology, and parameters were analyzed. A. Representative morphology of epididymal sperm collected from a pair of 6-week-old (a) and a pair of 3-month-old control and cKO mice. The inserts showed the zoom-in images. Sperm from the 6-week-old mice appeared to be normal. However, less sperm density was observed in the 3-month-old cKO mouse, and many of them were not normal. B. Sperm parameters of the mice at the indicated ages. a: sperm count; b: percentage of normal morphology; c: percentage of motile sperm; d: sperm motility. *P<0.05; **P< 0.01; *** P<0.001. M: month. n≥6.

### Disruption of *Cops5* gene in Sertoli cells resulted in progressive testis atrophy and spermatogenesis defects

Sperm are made in the testis. Given the progressively reduced male fertility, sperm count and motility of the cKO mice and testes of the control and the cKO mice were examined. There was no significant difference in testis/body weight when the mice were 6-week-old and 2-month-old. However, from 3 months old, testis weight and testis/body weight were significantly lower in the cKO mice, and the reduction was age dependent. Interestingly, even though no significant difference was found in the body weight between the control and cKO mice when the mice before 3-months-old, the cKO mice had a greater body weight when the mice were 3-month-old and older (**Figure 3**).

**Figure 3.**
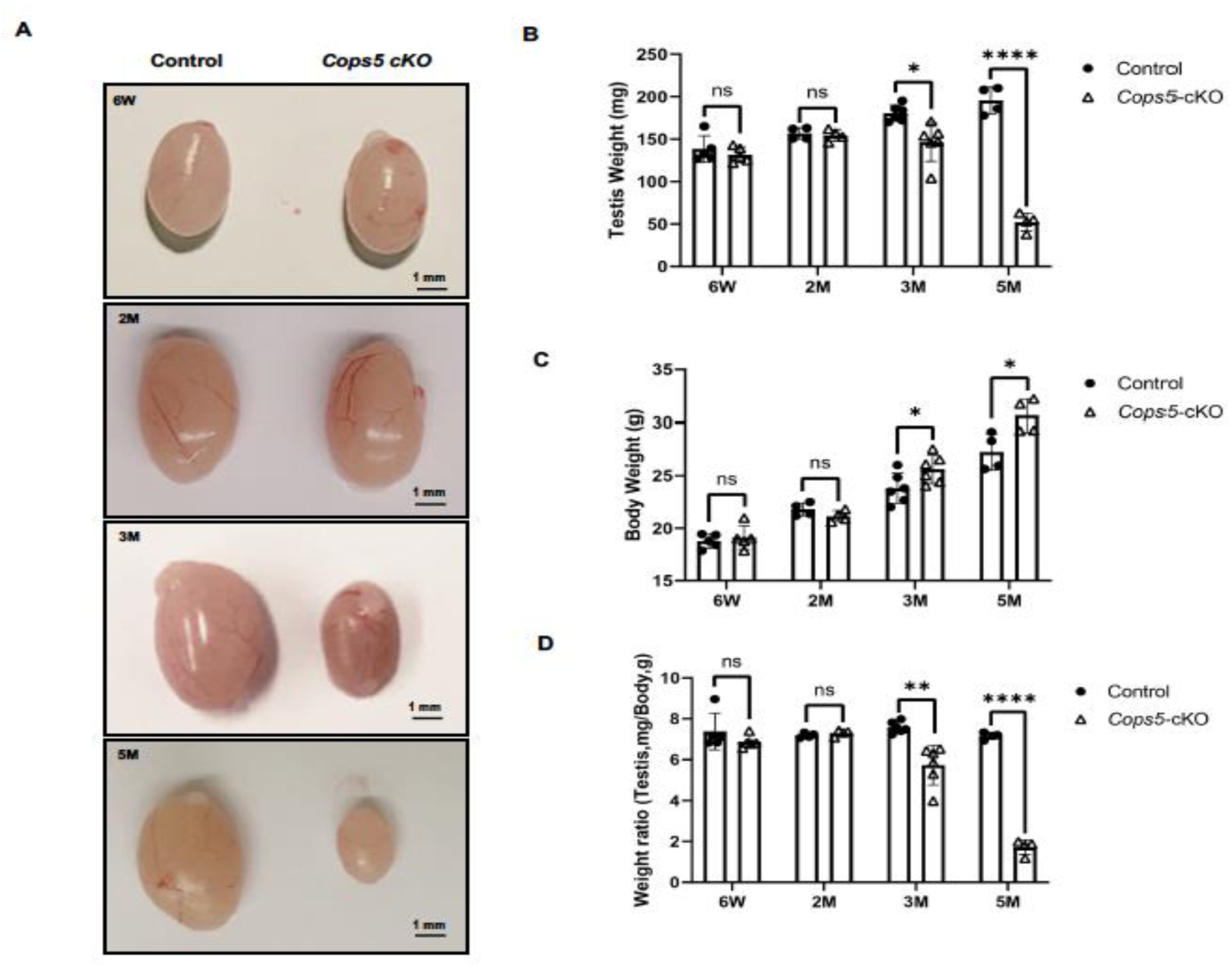
The testes in the cKO mice undergo progressive testis atrophy with age. A. Representative testis images of the control and cKO mice. No difference was observed between the control and cKO mice at 6-week-old and 2-month-old. The testes became smaller after 3-month-old. B. Testis weight of control and cKO mice. No significant difference was found between the control and cKO at 6-week-old and 2-month-old mice. The testis weight was significantly lower in the 3-month-old cKO mice and after. C. Body weight of control and cKO mice. No significant difference was found between the control and cKO at 6-week-old and 2-month-old mice. The testis weight was significantly higher in the 3-month-old cKO mice and after. D. Testis/body weighty of control and cKO mice. No significant difference was found between the control and cKO at 6-week-old and 2-month-old mice. The testis weight was significantly lower in the 3-month-old cKO mice and after. *P<0.05; **P< 0.01; *** P<0.001. M: month. n≥6.

Given that testis atrophy was observed in an age-dependent manner, testis histology of the mice at these ages was further examined by H&E staining. No difference was observed between the control and cKO mice when 6-week-old and 2-month-old mice were analyzed (**Supplemental Figure 2**, **Figure 4Aa, b**). At 5-months-old, severely disrupted seminiferous tubule architecture was observed in the cKO mice, including vacuolation, a substantial depletion of germ cells, and very few spermatocytes or elongated spermatids. However, the seminiferous tubules of the age-matched control mice showed well-organized layers of germ cells and abundant sperm flagella in their lumina (**Figure 4Ae, f**). Further analysis demonstrated that the diameter and cross-sectional area of the seminiferous tubules were significantly reduced in both 2- and 5-month-old *Cops5* cKO mice compared to controls, confirming that the loss of *Cops5* in Sertoli cells leads to testicular atrophy, which aggravates with advancing age (**Figure 4Ba, b**). Cauda epididymis histology was also examined in these 2-month-old and 5-month-old control and cKO mice. In all control mice analyzed, the lumen of the cauda epididymis was filled with sperm **(Figure 4Ac, g**). In the 2-month-old *Cops5* cKO mice, the lumen was also filled with sperm (**Figure 4Ad**). However, in the 5-month-old *Cops5* cKO mice, the lumen was almost empty (**Figure 4Ah**).

**Figure 4.**
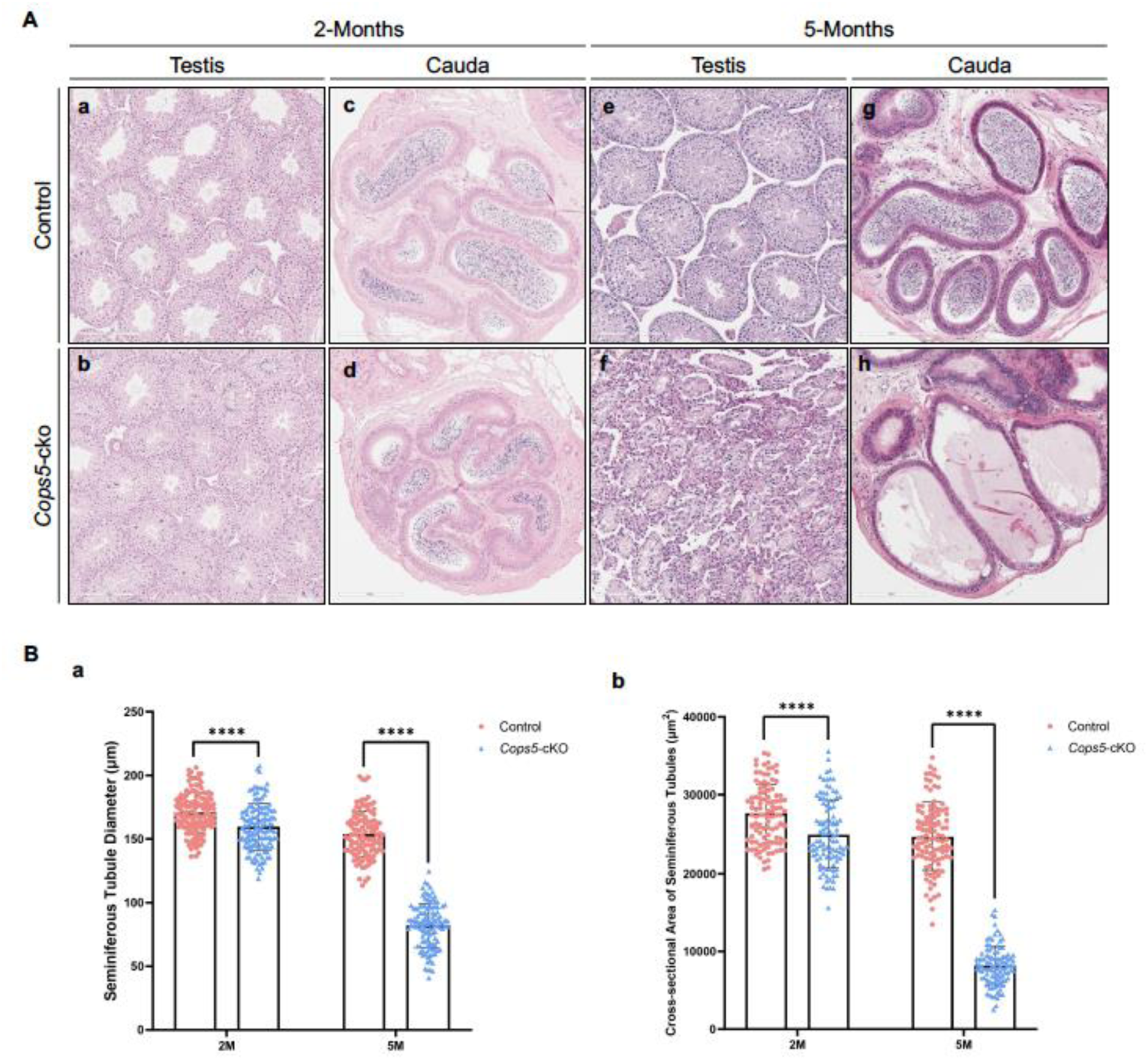
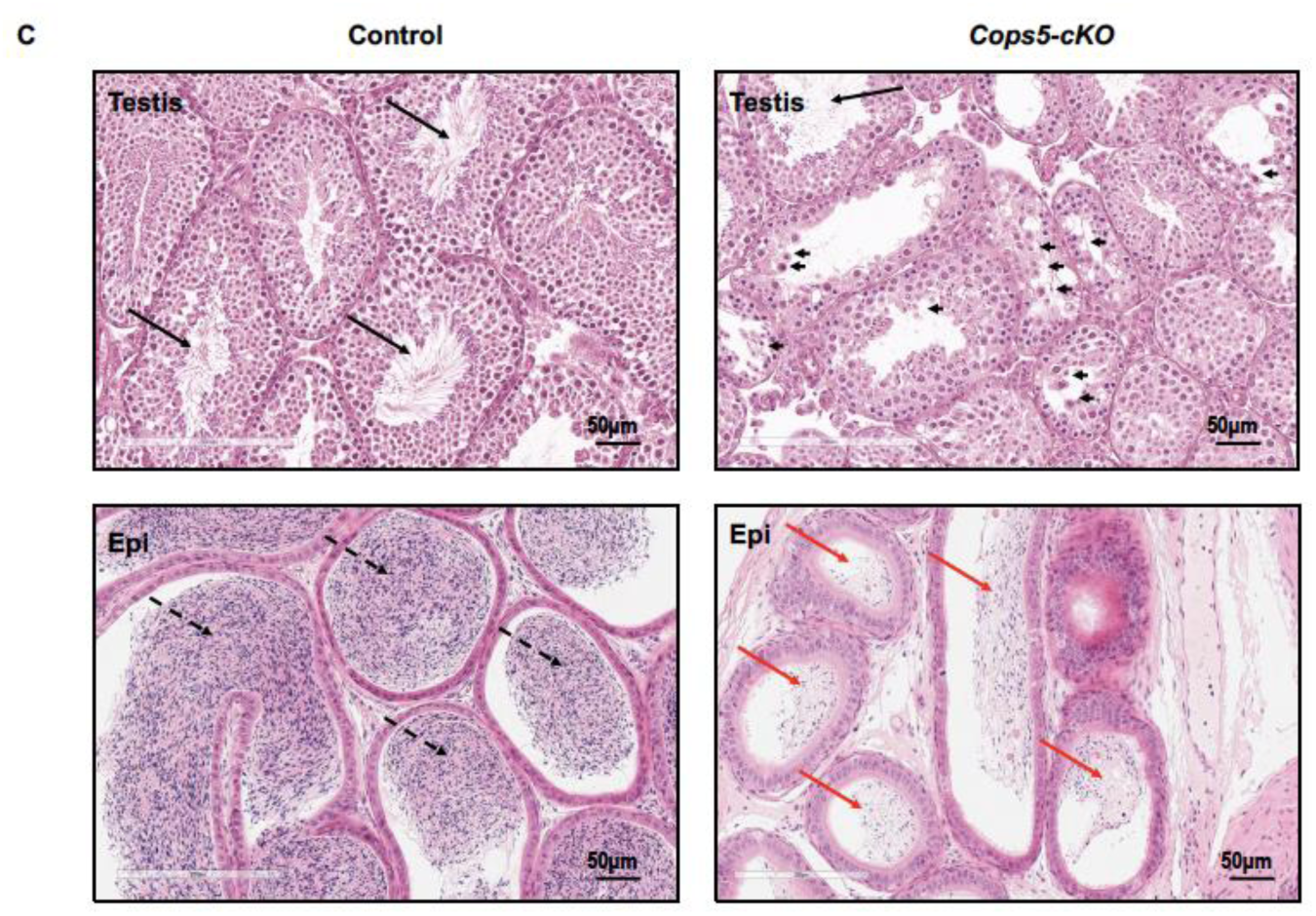
Targeting *Cops5* in Sertoli cells results in progressively disruptive spermatogenesis. A. H&E staining of testis and cauda epididymis of 2-month and 5-month-old control and *Cops5* cKO mice. At 2-month-old mice, no difference was found in the control and *Cops5* cKO testis (a and b) and cauda epididymis (c and d). At 5-month-old mice, the control mice showed normal testis histology (e), but the *Cops5* cKO mice started showing dramatic defects in seminiferous tubules (f). The cauda in the control mice was filled with sperm (g). However, sperm could be hardly seen in the *Cops5* cKO mice (h). B. Significant reduced seminiferous tubule diameter and cross-sectional area of the in the cKO mice. The diameter of the seminiferous tubules was measured, and the cross-sectional area was calculated. it was found that a significantly reduced diameter and cross-sectional area in both 2-month-old and 5-month-old *Cops5* cKO mice. a: seminiferous tubule diameter; b: cross-sectional area. C. Histology of testis and cauda epididymis of the 3-month-old control and *Cops5* cKO cKO mice. The seminiferous tubules were largely disrupted in the *Cops5* cKO mice. The black arrows point to the sperm in the seminiferous tubule lumen. The arrow heads point to the degenerated germ cells in the seminiferous tubule lumen in the *Cops5* cKO mice; the dashed arrows point to the sperm in the cauda epididymis lumen of the control mice; the red arrow bars point to the lumen of cauda epididymis of the *Cops5* cKO.

Testis atrophy was observed in 3-month-old *Cops5* cKO mice. We therefore examined histology of 3-month-old control and *Cops5* cKO mice. The control mouse testis showed normal architecture, and sperm were observed in the lumen of the seminiferous tubules and cauda epididymis (**Supplemental Figure 3, Figure 4C**) started showing defects in the cKO mice. However, significantly reduced sperm was found in the lumen of seminiferous tubules and cauda epididymis in the *Cops5* cKO mice (**Figure 4C**). A large number of degenerated germ cells were present in the seminiferous tubule lumen (**Figure 4C).**

### Disrupted Sertoli cell distribution in the seminiferous tubules and subcellular localization of key protein in Sertoli cells when *Cops5* is disrupted in Sertoli cells

Sertoli cells anchor to the basement membrane on their basal side, which mediate material exchange with the blood (e.g., uptake of nutrients and hormonal signals like testosterone and FSH) and hosting spermatogonial stem cells and primary spermatocytes. Heir apical side projects into the tubule lumen, supporting advanced germ cell development (e.g., spermatids) and releasing sperm. We further stained the testicular tissues with WT1, a Sertoli cell marker expressed consistently from embryonic stages to adulthood ^[31]^. In adult testes, WT1 is specifically localized in the nuclei of Sertoli cells and is a key indicator for their function and testicular development. In both 2-month and 5-month-old control mice, all Sertoli cell nuclei showed strong and orderly WT1 staining along the basement membrane (**Figure 5Aa, c**). However, in *Cops5* cKO mice, even though WT1 signal was the same as the control mice (**Figure 5Ab**), WT1 was partially localized to the cytoplasm in the 5-month-old mice. Additionally, some Sertoli cells detached from the basement membrane and moved toward the lumen while others became sparse and irregularly arranged, and some lost directional alignment (**Figure 5Ad**).

**Figure 5.**
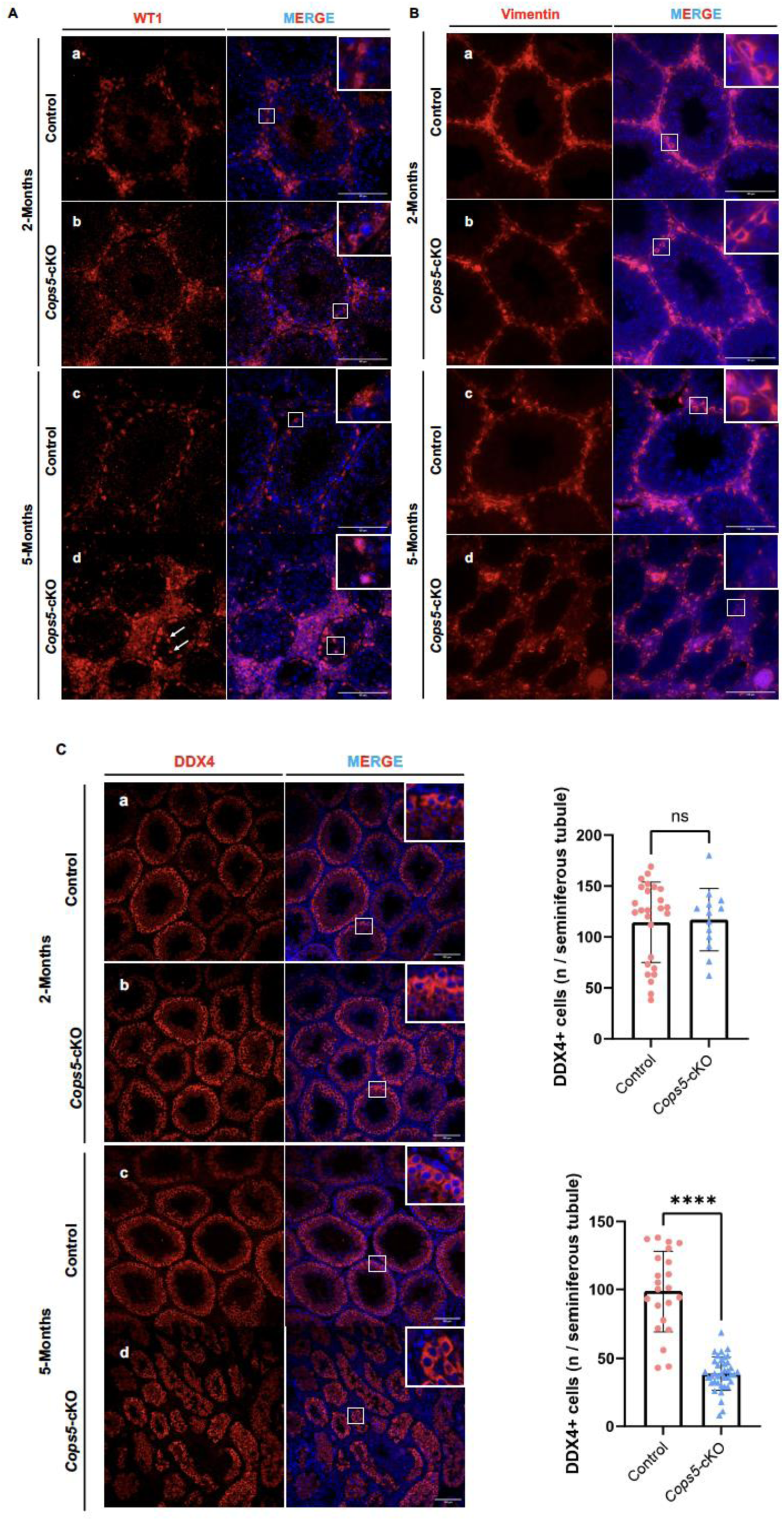
Disrupted Sertoli cell distribution, subcellular localization of key protein and loss of germ cells in the absence of COPS5 in Sertoli cells. A. Immunofluorescence result of WT1 in control and *Cops5* cKO mice. In 2-month-old control and *Cops5* cKO mice, as well as 5-month-old control mice, WT1 signal is in the nuclei of Sertoli cells along the basement membrane. However, in 5-month-old *Cops5* cKO mice, some WT1 signals were present in cytoplasm with intense expression. In some Sertoli cells, although WT1 was in the nuclei, but the protein detached from the basement membrane and moved toward the lumen; others became sparse and irregularly arranged, and some lost directional alignment. B. Immunofluorescence result of vimentin in control and *Cops5* cKO mice. In 2-month-old control and *Cops5* cKO mice, as well as 5-month-old control mice, strong vimentin signal was observed in the perinuclear region, and vimentin-positive extensions, projecting toward the lumen. However, in 5-month-old *Cops5* cKO mice, some cells maintained a similar pattern. In many cells, only weak, dot-like vimentin signals were observed. C. Immunofluorescence result of DDX4 in control and *Cops5* cKO mice. a. Representative images of DDX4 staining in 2 and 5-month-old control and *Cops5* cKO mice. Notice that the DDX4 staining pattern was the same in all the samples. However, in the 5-month-old *Cops5* cKO mice, the number of DDX4 positive cells was dramatically reduced. b. Statistic analysis of DDX4 positive cells/seminiferous tubule between the control and *Cops5* cKO mice. There was no significant difference between 2-month-old control and *Cops5* cKO mice. However, in 5-month-old mice, the *Cops5* cKO mice had significantly reduced DDX4 positive cells/seminiferous tubule.

Another Sertoli cell marker, Vimentin, was also examined ^[32]^. In both 2-month and 5-month-old control mice, strong vimentin signal was observed in the perinuclear region and vimentin-positive extensions, projecting toward the lumen (**Figure 5Ba, c**). The same pattern was seen in the 2-month-old *Cops5* cKO mice (**Figure 5Bb**). However, in the 5-month-old *Cops5* cKO mice, the pattern was largely changed. Even though some cells maintained a similar pattern. In many cells, only weak, dot-like vimentin signals were observed (**Figure 5Bb**).

### Disruption of *Cops5* in Sertoli cells resulted in progressive germ cell loss in the seminiferous tubules

To explore the effect of disrupting *Cops5* specifically in Sertoli cells on germ cells, we used an anti-DDX4 (a germ cell marker) ^[33]^ antibody for immunofluorescence staining on testicular tissues in 2-month and 5-month-old *Cops5* cKO and control mice. In 2-month-old mice, there was no significant difference in germ cell number between the control mice and the *Cops5* cKO and control mice. However, in 5-month-old mice, there was a significant decrease in the average number of germ cells per seminiferous tubule in the *Cops5* cKO mice (**Figure 5C**).

### Disruption of *Cops5* gene in Sertoli cells also affects blood-testis barrier and Gap junction

Sertoli cells display obvious morphological, structural, and functional polarity. This polarity is crucial for forming the blood-testis barrier (BTB) and for supporting the organized development and release of sperm. The specific localization of proteins and organelles within Sertoli cells maintains this polarity. Tight junction (TJ) proteins, key components of the BTB, are specifically localized to a particular region of the lateral cell membrane. ZO-1, a tight junction protein, is a key marker for Sertoli cell function and BTB integrity ^[34–35]^, and its mislocalization directly indicates lost polarity and BTB impairment. In 2 and 5-month-old control mice and 2-month-old *Cops5* cKO mice, ZO-1 formed continuous, fence-like lines around the base of the seminiferous tubules, clearly separating the basal and adluminal compartments. In contrast, in 5-month-old *Cops5* cKO testes, the ZO-1 signal was discontinuous, fainted, and punctate (**Figure 6A**), indicating defective tight junctions and a disrupted BTB structure due to the loss of COPS5 protein.

**Figure 6.**
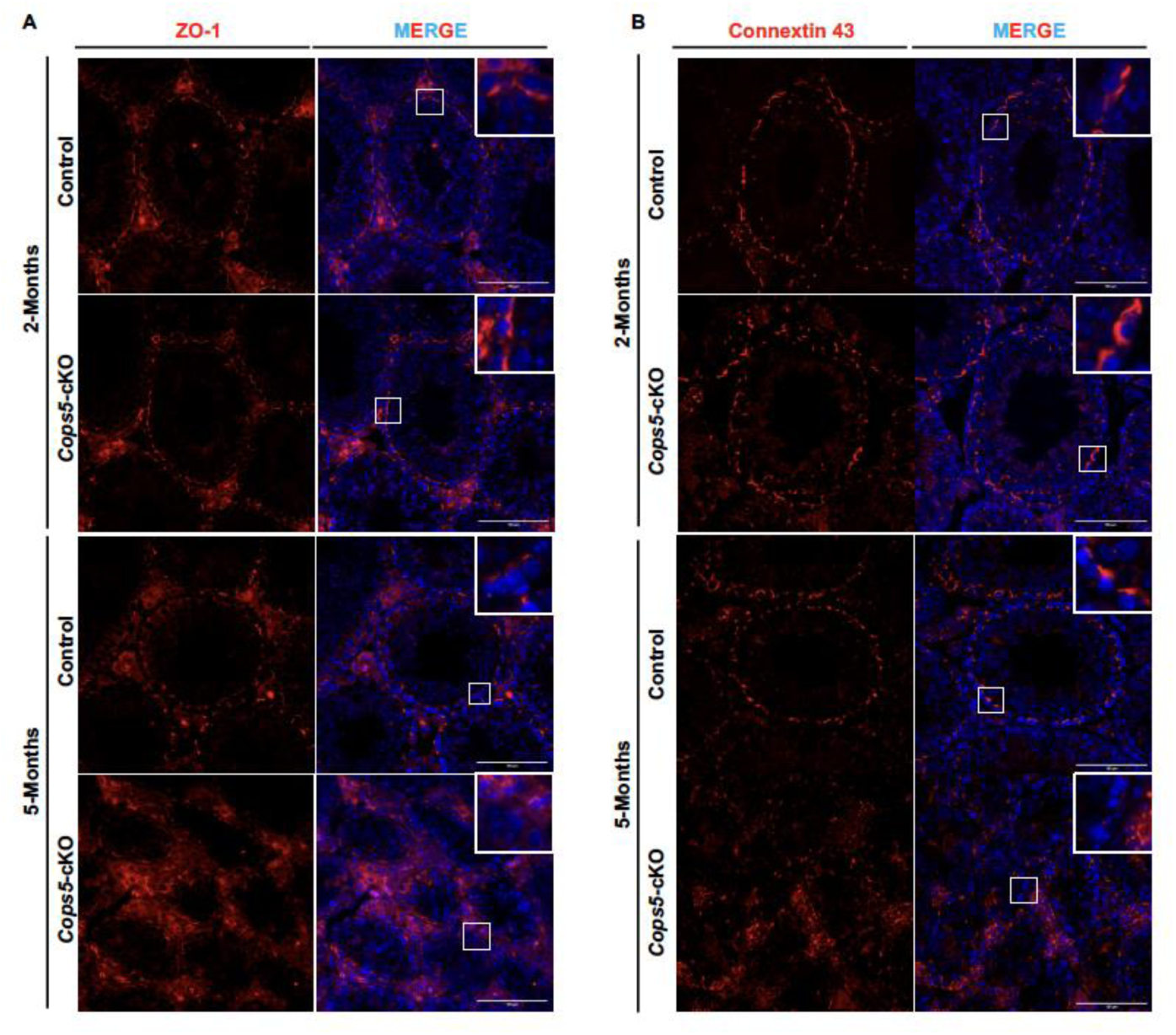
Disruption of *Cops5* gene in Sertoli cells affects blood-testis barrier and Gap junction. A. Analysis of ZO-1 staining pattern. ZO-1, a tight junction protein, and a key marker for Sertoli cell function and BTB integrity was analyzed by Immunofluorescence staining. In 2 and 5-month-old control and 2-month-old *Cops5* cKO mice, ZO-1 formed continuous, fence-like lines around the base of the seminiferous tubules, clearly separating the basal and adluminal compartments (A). In contrast, in 5-month-old *Cops5* cKO mice, the ZO-1 signal was discontinuous, faint, and punctate, indicating a disrupted BTB structure. B. Analysis of Connexin 43 staining pattern. Connexin 43 (Cx43) is a key component of gap junction. In 2 and 5-month-old control and 2-month-old *Cops5* cKO mice, Cx43 appeared as distinct, discrete punctate spots at cell-cell contacts near the basement membrane. This signal was greatly reduced or lost in the 5-month-old *Cops5* cKO mice.

Gap junctions (GJ) are channels between Sertoli cells and between Sertoli cells and spermatogonia. They allow direct transfer of ions, second messengers (like cAMP and Ca²⁺), and small metabolites, acting as vital communication hubs for maintaining polarity and the cellular environment. Connexin 43 (Cx43) is the primary junction protein in the testis ^[36]^. In 2 and 5-month-old control mice, and 2-month-old *Cops5* cKO mice, Cx43 appeared as distinct, discrete punctate spots at cell-cell contacts near the basement membrane. This signal was greatly reduced or lost in 5-month-old *Cops5* cKO testes (**Figure 6B**). This abnormal expression of Cx43 suggests that *Cops5* deficiency disrupts cell communication, likely impairing BTB function, disturbing the stem cell niche, and causing spermatogenesis defects.

Collectively, the proper function of both ZO-1 and Cx43 is required to maintain Sertoli cell polarity and spermatogenesis. The defects in both proteins upon *Cops5* deletion demonstrate that COPS5 is essential for preserving BTB integrity and Sertoli cell polarity.

## Discussion

The mouse testis is a highly organized organ where spermatogenesis occurs within the seminiferous tubules. This process is critically supported and regulated by a consortium of somatic cells, which create the necessary structural, nutritional, and immunological environment ^[7,37–38]^. The primary somatic cell types within the testis include: Sertoli cells, Leydig cells, PTM cells and so on ^[39]^. Sertoli cells are the cornerstone of the seminiferous epithelium, providing structural, nutritional, and immunological support for developing germ cells. Their functions are multifaceted, including the formation and dynamic regulation of the BTB, the secretion of essential factors and nutrients, the phagocytosis of apoptotic germ cells and residual bodies, and the release of signals that orchestrate the complex process of spermatogenesis ^[22,39–41]^. The integrity of Sertoli cell function is absolutely prerequisite for the completion of meiosis and spermiogenesis ^[42]^. In this study, we investigated the role of COPS5 in Sertoli cells by using the *Amhr2-Cre* to disrupt the *Cops5* gene. *Amhr2* gene encodes the anti-Mullerian hormone (AMH) receptor 2 ^[43–44]^. It is largely expressed in the Sertoli cells. Therefore, the *Amhr2-Cre* mice were used to disrupt genes in Sertoli cells.

Similar to the reproductive phenotypes observed in germ cell ^[19]^ and smooth muscle cell-specific ^[20]^ *Cops5* knockout mice, Sertoli cell-specific ablation of *Cops5* led to smaller testes, morphological abnormalities and reduced sperm count in the epididymis, lower sperm motility, and infertility in male mice. This provides additional evidence for the essential role of COPS5 in the male reproductive system, highlighting its necessity not merely in germ cells and testicular smooth muscle cells, but equally in Sertoli cells. The phenotype confirms that COPS5 is intrinsically necessary for Sertoli cells to sustain and promote germ cell development during spermatogenesis. Strikingly, deletion of *Cops5* in Sertoli cells disrupted the expression pattern of the Sertoli marker WT1 and Vimentin, but not the germ cell marker DDX4, indicating that the disruption is Sertoli cell specific.

It is interesting, the reproductive phenotype was not observed in younger mice (2-month-old), but from 3-month-old mice, and the phenotype became more significant in older mice, this might be caused by the late expression of Cre recombinase driven by the *Amhr2* promoter. Even though the endogenous *Amhr2* gene is expressed during embryonic stage ^[44]^. The high cre recombinase expression in the *Amhr2-cre* transgenic mice is postnatal 2-3 weeks ^[45–46]^. Therefore, the *Cops5* gene disruption deficiency in Sertoli cells might not be high enough, and no phenotype was observed before 3-month-old. After the mice are 3-month-old, the cre activity is high enough to disruption *Cops5* gene in Sertoli cells, which result finally result in Sertoli cell failure.

A fundamental aspect of Sertoli cell function is the establishment and maintenance of cell polarity ^[40,47–48]^. Sertoli cells display a highly polarized architecture with distinct apical, lateral, and basal membrane domains. This polarity is crucial for the directional secretion of nutrients, the formation of the BTB at the basal compartment, and the proper release of spermatozoa at the apical side (spermiation) ^[48]^. Disruption of Sertoli cell polarity is a well-established mechanism leading to spermatogenic failure ^[40,47–48]^. In our cKO mice, we further detected aberrant expression of the tight junction protein ZO-1 and the gap junction integral membrane protein Connexin 43 (CX43). ZO-1 is a crucial cytoplasmic scaffolding protein that acts as a “bridge,” linking transmembrane proteins (such as Occludin and Claudins) to the intracellular actin cytoskeleton ^[34]^. By anchoring these transmembrane proteins to actin, ZO-1 facilitates the assembly, stabilization, and maintenance of the structural integrity of tight junctions, forming a continuous “fence”-like structure ^[49–50]^. Concurrently, ZO-1 serves as a signaling platform involved in regulating the dynamics of the Blood-Testis Barrier (BTB). It can recruit signaling molecules and enzymes in response to external stimuli (e.g., hormones, cytokines), thereby coordinating the opening and closing of the BTB to permit the migration of germ cells ^[35,51–52]^. Connexin 43 (Cx43) is a transmembrane protein that constitutes gap junctions ^[36,53–55]^. Six Cx43 proteins assemble to form a connexon (hemichannel) on the Sertoli cell membrane, which then docks with a connexon on the membrane of an adjacent cell (another Sertoli cell or a germ cell) to form a complete hydrophilic channel. These channels allow for the direct intercellular exchange of ions (e.g., Ca²⁺), second messengers (e.g., cAMP, IP₃), small metabolites (<1 kDa), and microRNAs. They collectively contribute to maintaining BTB function and the intraluminal environment. The channels between Sertoli cells and germ cells (particularly spermatogonia) are vital for transmitting key signaling molecules that regulate spermatogenesis and for sustaining the spermatogonial stem cell niche ^[56–57]^. This loss of polarity directly explains the observed testicular atrophy and germ cell loss: a compromised BTB fails to create a protected microenvironment for meiosis, and defective nutritional support and signaling from apically disordered Sertoli cells lead to germ cell apoptosis and failed spermiation ^[58]^.

Even though it has been reported that *Amhr2-Cre* is also expressed in other cell types in the testis ^[45,59]^, we believe that the phenotype described here is largely due to the disruption of the *Cops5* gene in Sertoli cells. We previously disrupted *Cops5* gene in male germ cells using *Stra8-iCre* and Smooth muscle cells with *MYH11-Cre* ^[19–20]^, the phenotypes were different. In the germ cell specific cKO mice, the Sertoli cells were normal, but germ cell underwent apoptosis ^[19]^. When *Cops5* gene was disrupted through *MYH11-Cre* mice, the cKO mice showed growth retardation, a variety of developmental and reproductive disorders, including failure of development of reproductive organs, particularly dramatic impairment of the endocrine system associated with testicular functions, including a marked reduction in serum levels of gonadotropins ^[20]^. These phenotypes are different from what we observed in the current model.

The 3-month-old cKO mice showed increased body weight, and this becomes more significant in the 5-month-old cKO mice. The Sertoli cells also conduct endocrine function. The change in body weight might be caused by disruption of this function In Sertoli cells. In addition, Cre activity is also expressed other than Sertoli cells, including Leydig cells. Disruption of *Cops5* in these cells might also cause increased body.

In conclusion, our findings firmly establish COPS5 as an essential regulator in Sertoli cells, the pivotal somatic component of the testis. The severe spermatogenic defect resulting from its ablation underscores that its function is non-compensable and critical for male fertility. This study, combined with previous work on its roles in germ cells and peritubular myoid cells, paints a picture of COPS5 as a pleiotropic master regulator that is indispensable across multiple testicular cell types. These results significantly advance our understanding of male infertility and highlight the paramount importance of investigating crucial regulators like COPS5 within the context of specific somatic cells in the reproductive system. Future research should focus on delineating the precise molecular pathways through which it governs these phenotype. Such efforts could unveil novel therapeutic targets for the treatment of idiopathic male infertility.

## Data Availability

The data is available from the corresponding author upon reasonable request.

## Author Contribution

Designing research studies: Zhibing Zhang.

Conducting experiments: Changmin Niu, Tao Li, Wei Li, Yi Tian Yap, Qian Huang, Opeyemi Dhikhirullahi, Ava Miciuda, Eva Faddoul,

Acquiring data: Changmin Niu, Tao Li, Wei Li, Yi Tian Yap, Qian Huang, Opeyemi Dhikhirullahi.

Analyzing data: Lei Jiang, Shizheng Song, Michael D Griswold, Zhibing Zhang.

Managing project: Zhibing Zhang

The authorship order was assigned depending on the contributions.

## Acknowledgments

This research was supported by the Wayne State University Start up Fund and NIH RO1 Awards (HD105944, HD114311).

## Figure legends

**Supplemental Figure 1.**
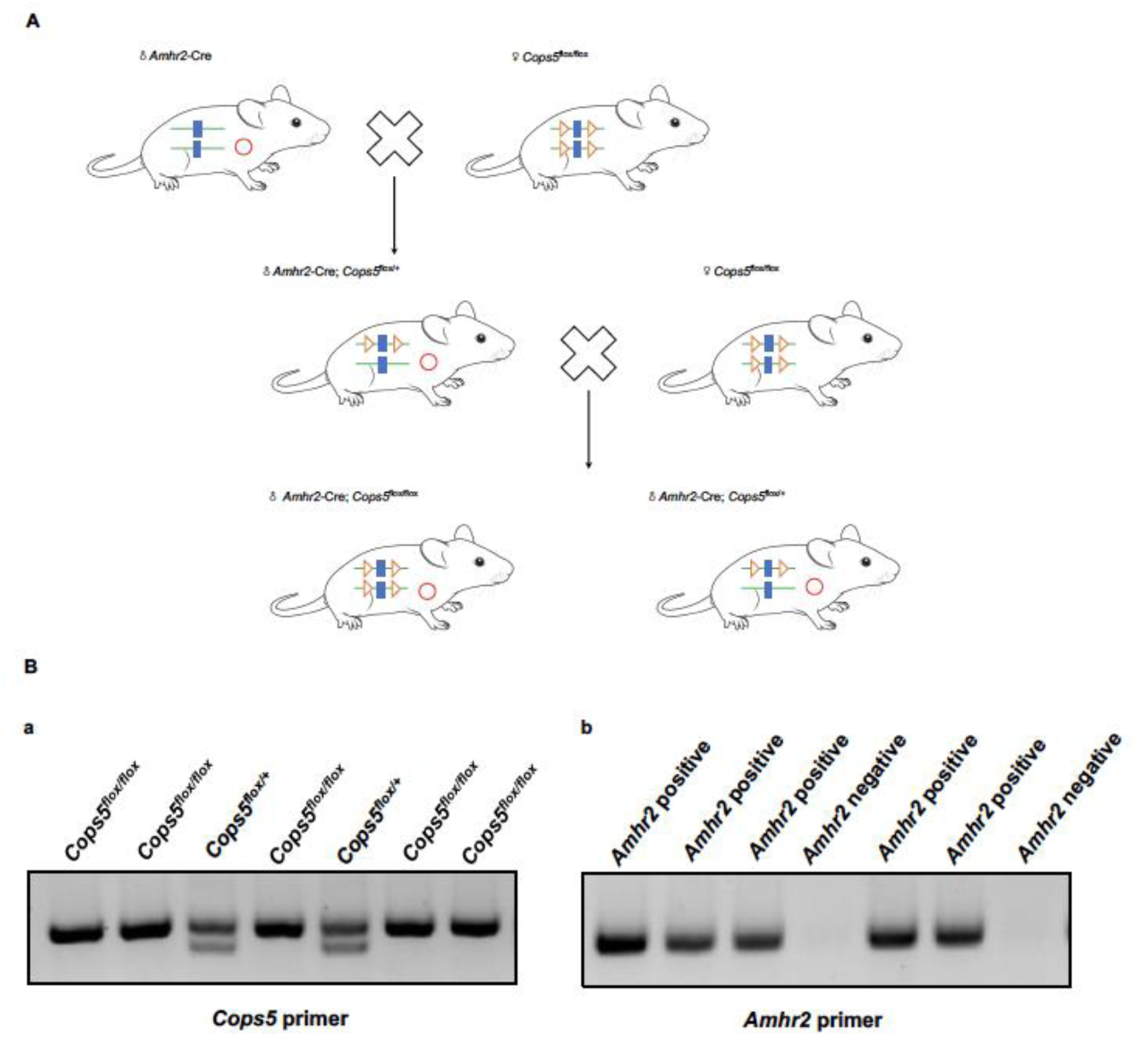
Generation of Sertoli cell-specific Cops5 KO mice. A. Breeding strategy to generate Cops5 cKO mice; B. Representative PCR result for mouse genotyping.

**Supplemental Figure 2.**
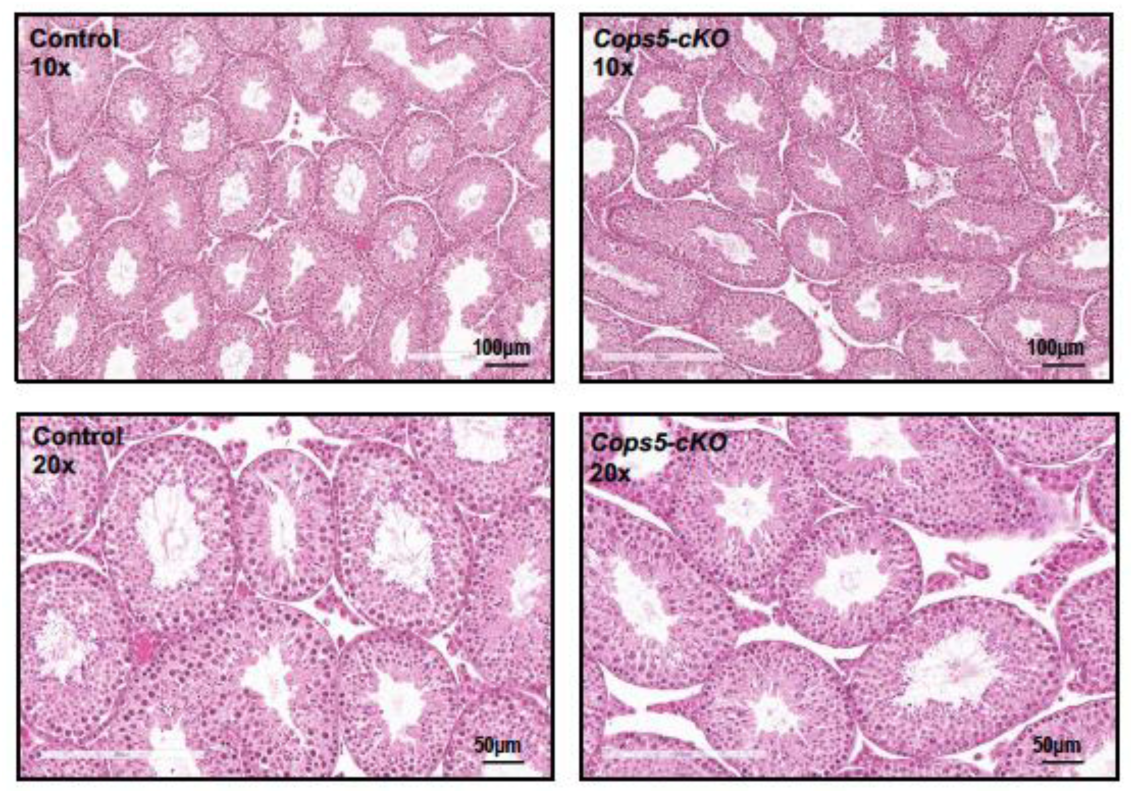
Testis histology of 6-week-pld control and *Cops5* cKO mice. Representative images of H&E staining of testis from 6-week-old control and *Cops5* cKO mice. No difference was observed between the two genotypes.

**Supplemental Figure 3.**
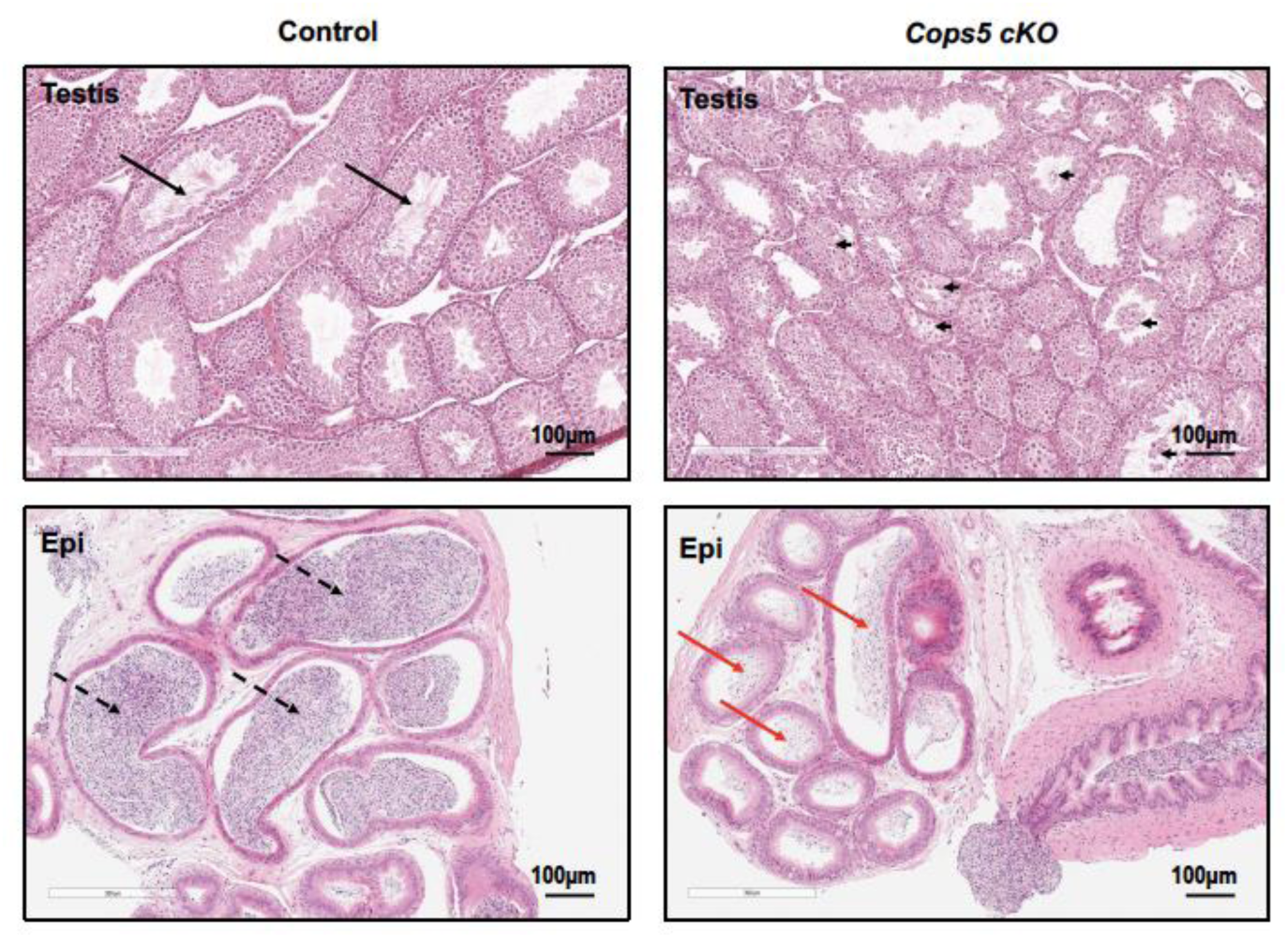
Testis histology of 3-month-old control and *Cops5* cKO mice at low magnification. The black arrows point to the sperm in the seminiferous tubule lumen in the control mice. The dashed arrows point to the sperm in the cauda epididymis lumen of the control mice; the arrow heads point to the degenerated germ cells in the seminiferous tubule lumen in the *Cops5* cKO mice; the red arrows point to the few sperm in the cauda epididymis lumen of *Cops5* cKO *mice*.

